# Highly Effective Batch Effect Correction Method for RNA-seq Count Data

**DOI:** 10.1101/2024.05.02.592266

**Authors:** Xiaoyu Zhang

## Abstract

RNA sequencing (RNA-seq) has become a cornerstone in transcriptomics, offering detailed insights into gene expression across diverse biological conditions and sample types. However, RNA-seq data often suffer from batch effects, which are systematic non-biological differences that compromise data reliability and obscure true biological variation. To address these challenges, we introduce ComBat-ref, a refined method of batch effect correction that enhances the statistical power and reliability of differential expression analysis in RNA-seq data. Building on the foundations of ComBat-seq, ComBat-ref employs a negative binomial model to adjust count data but innovates by using a pooled dispersion parameter for entire batches and preserving count data for the reference batch. Our method demonstrated superior performance in both simulated environments and real datasets, such as the growth factor receptor network (GFRN) data and NASA GeneLab transcriptomic datasets, significantly improving sensitivity and specificity over existing methods. By effectively mitigating batch effects while maintaining high detection power, ComBat-ref proves to be a robust tool for enhancing the accuracy and interpretability of RNA-seq data analyses.

## Introduction

RNA sequencing (RNA-seq) has emerged as a pivotal technology in transcriptomics, providing unprecedented insights into gene expression profiles across different biological conditions and sample types. However, the reliability of RNA-seq data is frequently compromised by batch effects—systematic non-biological differences that arise during sample processing and sequencing across different batches. These batch effects can be on a similar scale or even larger than the effects of different biological conditions, significantly reducing the power of statistical analyses to detect differentially expressed genes.

The presence of batch effects in RNA-seq data has been widely acknowledged, and various strategies have been developed to mitigate their impact. ComBat [1] is a widely-used method that employs an empirical Bayes-based framework to correct both additive and multiplicative batch effects. SVASeq [2] and RUVSeq [3] are commonly used to model batch effects from unknown sources. Popular differential gene expression (DE) packages such as edgeR [4] and DESeq2 [5] also allow for the inclusion of batch as a covariate in the linear models to account for batch effects. ComBat-seq [6], which uses a general linear model (GLM) with a negative binomial distribution retains the integer count data and has shown better statistical power than ComBat and other previous methods. More recently, machine learning methods [7], [8] have also been proposed to model the data discrepancies among batches and eliminate batch effects from RNA-seq data.

Among the batch correction methods, ComBat-seq has the advantage of preserving the integer count matrix in adjusted data, making it suitable for subsequent differential gene expression analysis in edgeR and DESeq2. Furthermore, it has been shown to achieve higher statistical power in detecting DE genes than other methods when batches with different dispersion parameters are pooled together. This improved power is most likely due to its more accurate modeling of count data using negative binomial (gamma Poisson) distributions. However, ComBat-seq adjusted data still exhibit significantly lower power in DE analysis than data without batch effects, especially when using the false discovery rate (FDR; adjusted p-value) for statistical analysis, as recommended by software such as edgeR and DESeq2.

In this paper, we present a modified version of the batch effect adjustment method, named ComBat-ref, which models the RNA-seq count data using a negative binomial distribution similar to ComBat-seq, but with important changes in data adjustment. Firstly, ComBat-ref estimates a pooled (shrunk) dispersion for each batch and selects the batch with the minimum dispersion as the reference, to which the count data of other batches are adjusted. We demonstrate that ComBat-ref retains very high power, close to the level in data without batch effects, even when there is a large variance in the dispersion of batches. ComBat-ref performs especially well when using FDR for DE analysis, compared to other adjustment methods.

## Materials and Methods

Like ComBat-seq [6], we model the RNA-seq count data using a negative binomial distribution, with each batch potentially having different dispersions. Consider a gene *g* in batch *i* and sample *j*. Let *c*_*j*_ be the biological condition of sample *j* and 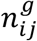 be the measured count. The count 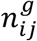 is modeled using a negative binomial distribution:

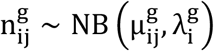

where 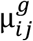 is the expected expression level of gene *g* in sample *j* and batch *i*, and 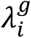 is the dispersion parameter for batch *i*. ComBat-seq estimates 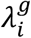 per gene and batch, then calculates an average dispersion per gene for data adjustment:

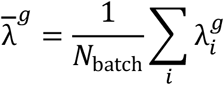

However, given that the number of samples per batch is usually small, we expect the estimation of 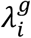 to be inaccurate with high variance, leading to high variance of 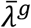 and reduced statistical power for adjusted data. In the new ComBat-ref method, we pool the gene count data in each batch and estimate a batch-wise dispersion *λ*_*i*_. The ComBat-ref method then selects the batch with the smallest dispersion as the reference batch. Without loss of generality, we assume the reference is batch 1. We noted that the alternative approach of ComBat [1], which uses a reference batch, was discussed in [9], but it did not employ a negative binomial model or select the reference based on dispersion.

We applied a generalized linear model (GLM) for the expected gene expression level 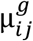 :

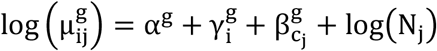

where *c*_*j*_ is the biological condition and *N*_*j*_ represents the library size of sample *j*· *α*^*g*^ denotes the global “background” expression of gene 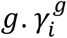 represents the effect of batch *i*, and 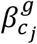 denotes the effects of the biological condition *c*_*j*_ on the logarithm expression level of gene *g*. The model parameters can be estimated using the GLM fit method implemented in edgeR[4], [10], or more computationally intensive MCMC based statistical methods [11]. Since the reference batch has the smallest estimated dispersion, it is desirable to keep its count data for down-stream DE analysis to achieve better statistical power. Therefore, we adjust RNA-seq count data in other batches toward the reference batch in the new ComBat-ref method.

### ComBat-ref Adjustment

Assuming reference batch 1 has the smallest dispersion *λ*_1_ among all batches, we compute the adjusted gene expression level 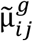 for batch *i* ≠ 1 and sample *j* as

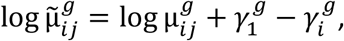

and set the adjusted dispersion simply as 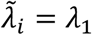. Then, similar to ComBat-seq, the adjusted count 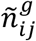is computed by matching the cumulative distribution function (CDF) of 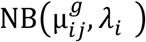 at 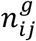 and CDF of adjusted distribution 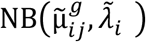 at 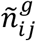. Care is taken to ensure that the adjusted count does not become infinity when the CDF is evaluated to be 1, and zero counts are always mapped to zeros.

Setting dispersion to *λ*_1_ enables high power in the subsequent statistical analysis of adjusted data, albeit with the potential cost of more false positives. It is often a preferred trade-off when samples from multiple batches are pooled for DE analysis. We noted in both simulations and real data that the ComBat-ref adjustment performed well with high sensitivity and FPR under control, especially when using adjusted p-values (FDR) in edgeR and DESeq2.

### Simulations

To evaluate the performance of the new ComBat-ref method, we utilized a similar simulation process as in ComBat-seq [6] to generate realistic RNA-seq count data to compare different batch correction (BC) methods. The count data were modeled with a negative binomial (gamma Poisson) distribution, assuming that different batches could affect the mean expression of genes and the dispersion of the distributions. The simulations assumed two biological conditions and two batches, with three samples for each combination (12 total samples). The simulation considers 500 genes, of which 50 are up-regulated and 50 are down-regulated with a mean fold change of 2.4. The batch effects are assumed to change the expression of genes by a fold factor (mean_FC) in one random batch, and increase the dispersion in batch 2 by a factor (disp_FC) over batch 1. We generated simulations for 16 experiments with varying batch effects on mean expression fold change (mean_FC = 1, 1.5, 2, 2.4) and dispersion fold change (disp_FC = 1, 2, 3, 4). There is no batch effect when mean_FC = 1 and disp_FC = 1. The gene-by-sample count matrices were generated using the polyester R package[12]. Results were averaged over 300 repeated simulated experiments with given parameters.

## Results

### Simulations

We first compared the performance of various batch correction (BC) methods in detecting differentially expressed (DE) genes, using edgeR [4] and the same threshold of an unadjusted p-value = 0.05 as in the original ComBat-seq paper [6]. Supplemental Figure 1 shows the edgeR results including those of the new ComBat-ref discussed in this paper. The results using the DESeq2 package were very similar. The simulation results of previous methods matched those in [6]. When there was no dispersion change between batches, as shown in the first column of Supplemental Figure 1Supplemental Figure 1, all methods achieved good true positive rates (TPR), while ComBat-ref demonstrated a slightly lower false positive rate (FPR) than the original ComBat-seq method. However, the new ComBat-ref achieved noticeably higher sensitivity than others in samples with larger dispersion fold changes (disp_FC), with TPR close to that of no batch effect. While the original ComBat-seq had slightly better TPR than the rest of the previous methods, the new ComBat-ref method had distinctively higher sensitivity, as the gaps became larger in the more challenging cases of increased disp_FC. The significantly improved sensitivity of the new ComBat-ref method did come at the cost of a higher type 1 error rate. In analyzing DE genes in samples plagued with batch effects, we often prefer the trade-off of higher power with potentially more false positives.

When we performed DE analysis using FDR and adjusted p-values, the advantage of ComBat-ref over other methods became more pronounced. The original ComBat-seq paper [6] did not include this analysis, possibly due to the very low power of all previous methods in difficult samples. As shown in Figure 1, ComBat-ref maintained the same level of power (TPR) as that of no batch effects, even in the most challenging case of mean_FC = 2.4 and disp_FC = 4. While using more stringent adjusted p-values and FDR ≤ 0.1 for statistical analysis, the other methods lost their power dramatically, with the discovery rate going down close to zero in the most challenging case. Not only did ComBat-ref maintain its near-ideal discovery rate, but it also had much smaller FPR than when using an unadjusted p-value. Based on the simulation results, we recommend using ComBat-ref to adjust for batch effects in RNA-seq count data and perform DE analysis using adjusted p-values for a better balance between type 1 and type 2 errors.

**Figure 1.**
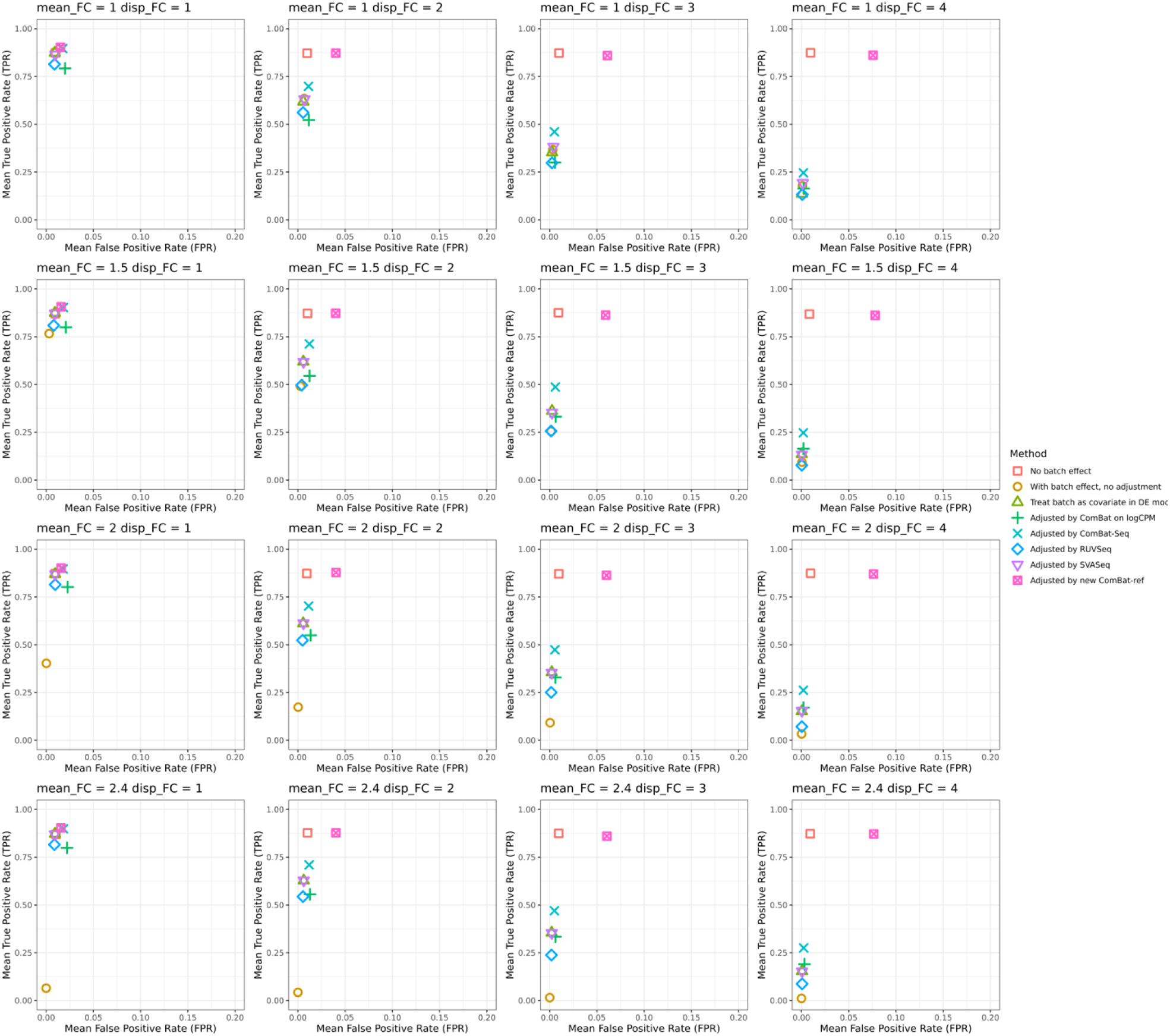
Simulation results using different levels of mean expression fold change (mean_FC = 1, 1.5, 2, 2.4) and dispersion fold change(disp_FC = 1, 2, 3,4). The DE genes for each method were identified using edgeR with an FDR≤0.1. As the mean expression and dispersion fold change increases between batches, all other methods except for ComBat-ref lost power significantly. ComBat-ref consistently maintains power for DE analysis, close to that of the data without batch effects, with controlled FPR, even in the most challenging experiment of mean_FC=2.4 and disp_FC=4.

### Applications to Real Data

We first tested the ComBat-ref method on the same growth factor receptor network (GFRN) data [13] used in the original ComBat-seq paper [6]. This dataset contains three batches, each introducing a specific GFRN oncogene to activate downstream pathway signals. Green fluorescent protein (GFP) controls were present in all batches. Batch 1 includes a total of 17 samples, five overexpressing HER2 and 12 GFP controls; batch 2 consists of 12 samples, six for EGFR and six controls; batch 3 comprises 18 samples, nine for KRAS and nine controls. The unadjusted data, shown in the first row of Figure 2, demonstrate strong batch effects where control samples and treated cells are grouped by batch. A successful batch adjustment should co-locate control samples while segregating treated cells from controls and each other.

**Figure 2.**
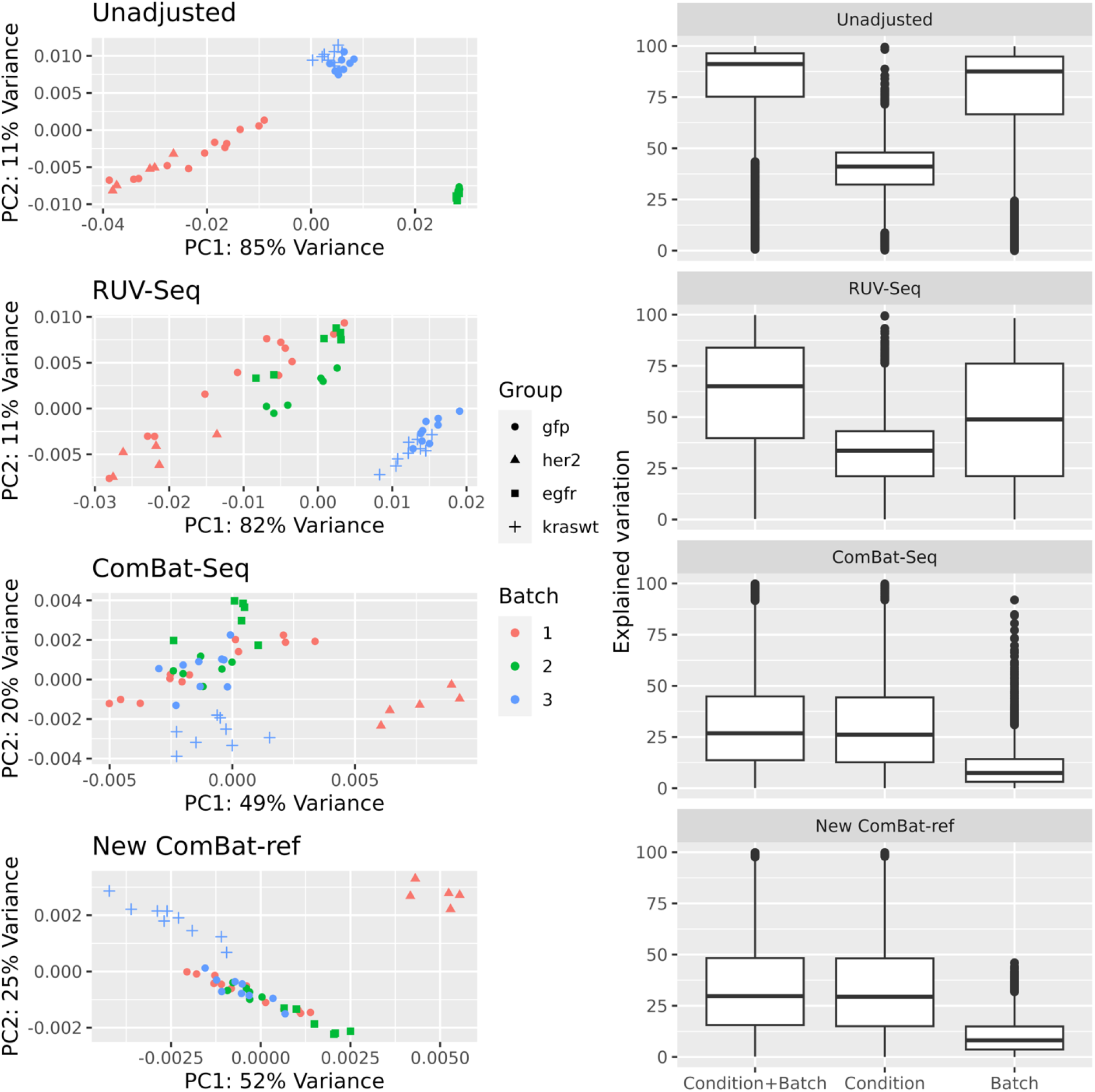
PCA plots of GFRN dataset for unadjusted data and data adjusted by RUV-Seq, ComBat-seq and the new ComBat-ref method respectively. The unadjusted data shows strong batch effects with all three batches are well separated. Both ComBat-seq and ComBat-ref can effectively bring the control (gfp) samples together and maintain biological signals from different treated samples. ComBat-ref did an even better job than ComBat-seq, verified by the quantitative measures in the text, also illustrated in the plots of explained variations by condition and batch.

We applied the new ComBat-ref method to the GFRN data and compared its results with those from RUV-Seq and ComBat-seq. Both ComBat-seq and ComBat-ref demonstrated effective batch adjustment, with control groups from different batches clustered together and treatment groups separated in the PCA plot. The boxplot showing the proportion of variation explained by condition and batch further demonstrated the effectiveness of batch correction using the ComBat-ref method. Compared to the original ComBat-seq method, ComBat-ref exhibited slightly higher variation for condition, while maintaining a similar reduction in variation explained by batch.

Further evaluation of clustering by group in the PCA plot was conducted using quantitative measures computed with the cluster.stats function from the fpc package [14], [15]. The unadjusted data had poor clustering evaluation scores by biological condition (the higher the better: Gamma = 0.19, Dunn = 0.78), where Gamma represents the correlation between distances and a 0-1 vector indicating different clusters; Dunn represents ratio of the minimum average inter-cluster dissimilarity to the maximum average intra-cluster dissimilarity. The data adjusted by the original ComBat-seq achieved much higher clustering scores (Gamma = 0.53, Dunn = 1.34). The scores for data adjusted by the new ComBat-ref method were even higher (Gamma = 0.66, Dunn = 1.60), indicating a superior batch effect correction outcome compared to ComBat-seq.

Analyses were then performed to identify DE genes between the treated samples and GFP controls pooled from the three batches, using the ComBat-ref method, along with other approaches studied in [6]. The results demonstrated that ComBat-ref outperformed ComBat-seq in statistical power in DE analysis, detecting more DE genes across all three conditions with an FDR ≤ 0.05 (Supplemental Table 1).

Furthermore, ComBat-ref appears to extract more biologically meaningful DE genes through batch correction. HER2, KRAS and EGFR were expected to over expressed in their respective samples. Like other methods, ComBat-ref successfully ranked HER2 and KRAS at the top of the DE gene list. However, detecting differential expression of EGFR proved more challenging due to batch effects. EGFR was not identified as DE in the unadjusted and several other adjusted datasets at an FDR ≤ 0.05. The success of the ComBat-seq method was demonstrated in [6] by its ability to detect EGFR as DE (FDR = 0.0016). ComBat also detected EGFR as DE (FDR = 0.0003). The new ComBat-ref method successfully detected EGFR as DE with a more statistically significant FDR=0.0001. The effect of ComBat-ref adjustment on the EGFR gene is shown in Supplemental Figure 2, illustrating a more distinct separation of its expression in EGFR and control samples than that of ComBat-seq, explaining the statistically significant DE results.

We also evaluated DE of the genes in the RAS signaling pathway for the KRAS sample, as done in [6]. Batch effect adjustment is expected to rank genes in the RAS signaling pathway among the top DE genes. In the top 1000 DE genes, ComBat-ref identified 30 genes in the RAS signaling pathway (Fisher’s exact test for enrichment, P = 0.0004), again outperforming ComBat (19 genes, P = 0.2) and ComBat-seq (24 genes, P = 0.02) (Supplemental Table 2).

Additionally, we applied the new ComBat-ref method to NASA GeneLab transcriptomic datasets [16], which include several mouse liver RNA-seq datasets from different space missions and library preparation technologies. These datasets contain multiple batch covariates, e.g.”mission” and “library preparation”, with researchers interested in differential expression of mouse liver genes in flight samples versus controls. The original study [16] found that batch adjustment by “library preparation” using ComBat was the most effective, followed by ComBat-seq. In batch correction with ComBat-ref, we treated them as three different batches rather than limiting them to a single batch factor.

We tested ComBat, ComBat-seq and ComBat-ref on a subset of NASA GeneLab transcriptomic datasets consisting of three batches: GLDS_137, GLDS_242 and GLDS_48 (Supplemental Table 3). These batches were selected due to their distinct separation in the PCA plot of unadjusted data (Supplemental Figure 3). We found that all three methods effectively remove most batch effects in the data set. GLDS_137 was identified to be the reference batch by ComBat-ref and its count data were not adjusted, thus the shape of its PCA plot remained similar to that in unadjusted data. While the data from the other two batches were adjusted to remove most batch effects, the grouping within each batch largely remained, leading to higher power in DE analysis. This result further demonstrated that the ComBat-ref batch correction method is applicable to RNA-seq count data from various sources and experiments.

## Discussion

We developed a simple and effective method, ComBat-ref, to correct batch effects in RNA-seq count data. Our simulations demonstrated that ComBat-ref greatly outperforms other methods in the statistical power of downstream differential expression analysis, achieving sensitivity (TPR) close to that of data without batch effects, while maintaining a controlled FDR. It also offers the same benefit as the ComBat-seq method in preserving integer count data, making it compatible with subsequent RNA-seq studies. When applied to the GFRN signal dataset, the ComBat-ref method showed superior performance in correcting batch effects and recovering biological signals compared to other methods, including ComBat-seq. We also demonstrated that ComBat-ref is generally applicable for correcting batch effects in other real datasets, such as the NASA GeneLab datasets.

Similar to ComBat-seq, ComBat-ref utilizes a GLM to model and adjust count data using the negative binomial distribution. However, to manage the additional variations introduced by the process, it estimates a per-gene expression effect for each batch but uses a pooled dispersion parameter for the entire batch. It preserves the count data for the reference batch and adjusts the other batches towards the reference. These seemingly minor but crucial modifications dramatically improve detection power in differential expression and better preserve biological signals in the adjusted data. It also addresses the issue of over-correction by ComBat-seq in data with little or no difference across batches.

Overall, our study has shown that ComBat-ref is an effective approach for adjusting general batched RNA-seq data. While it has been successfully applied to recover important biological signals from the GFRN dataset, it is imperative to test it on more datasets to evaluate its effectiveness further. The ComBat-ref method assumes a negative binomial distribution of count data with a pooled single dispersion for the entire batch, while the original ComBat-seq assumes a gene-wise dispersion parameter. Our test results in this paper have shown that our approach not only provides significantly higher test power but also comes with an elevated FPR. It would be interesting to see whether a more accurate estimation of dispersion could further enhance the effectiveness of batch correction, achieving similar power with fewer false positives.

## Reproducibility

Code to reproduce the results in this paper are available at https://github.com/xiaoyu12/Combat-ref

## Funding

National Institutes of Health (SC3GM122659)

### Conflict of interest statement

None declared.

## Supplemental Materials

**Supplemental Figure 1.**
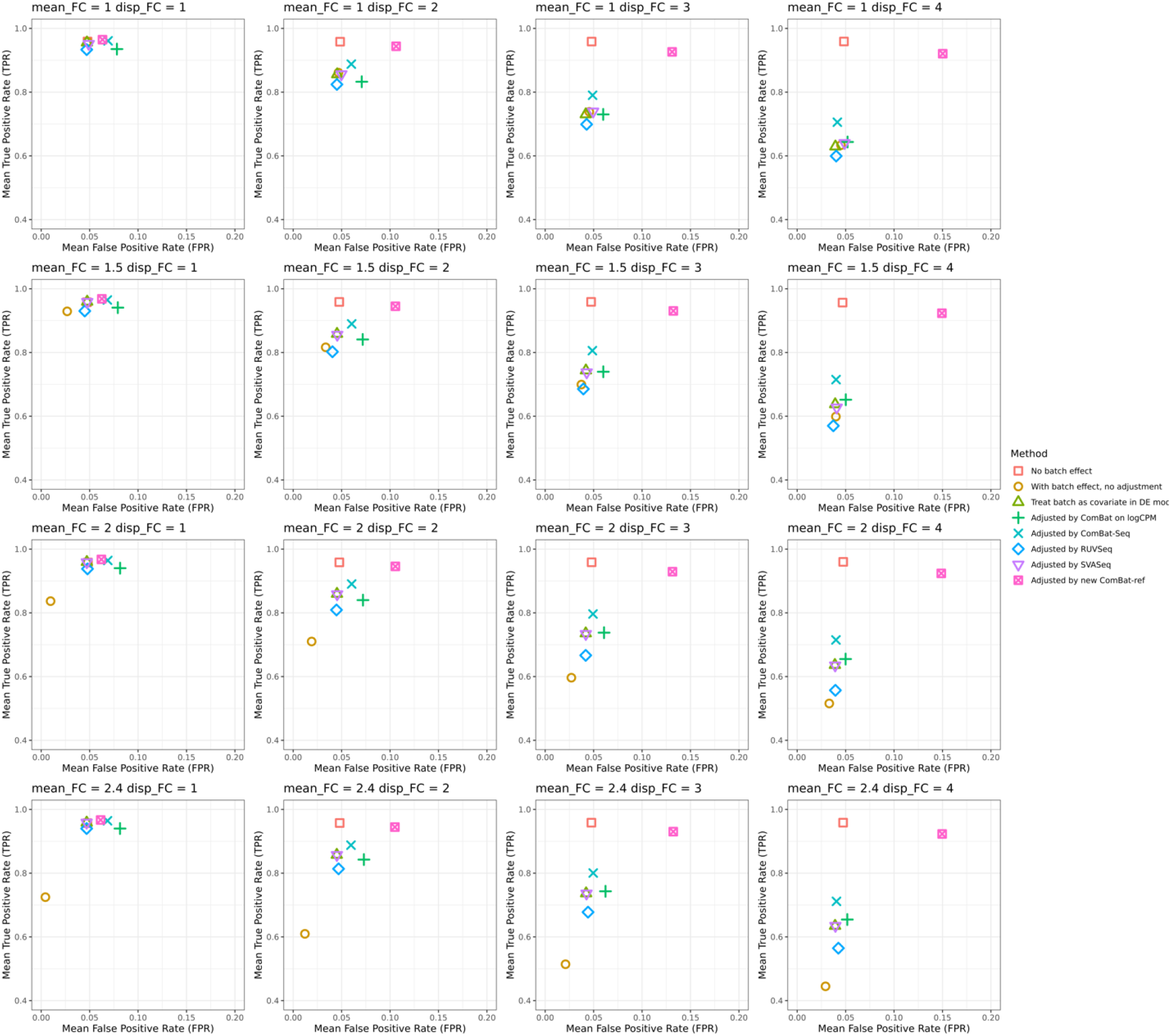
Simulation results using different levels of mean expression fold change (mean_FC = 1, 1.5, 2, 2.4) and dispersion fold change(disp_FC = 1, 2, 3,4). The DE genes for each method were identified using edgeR with an p-value⩽0.05. As the mean expression and dispersion fold change increases between batches, ComBat-ref outperforms all other batch correction methods, in term of power for DE analysis. However, ComBat-ref exhibits a higher false positive rate (FPR) than other methods when disp_FC increases. We recommend using an adjusted p-value (FDR) for DE analysis of ComBat-ref adjusted data in edgeR or DESeq2, which achieves better statistical results as shown in Figure 1.

**Supplemental Figure 2.**
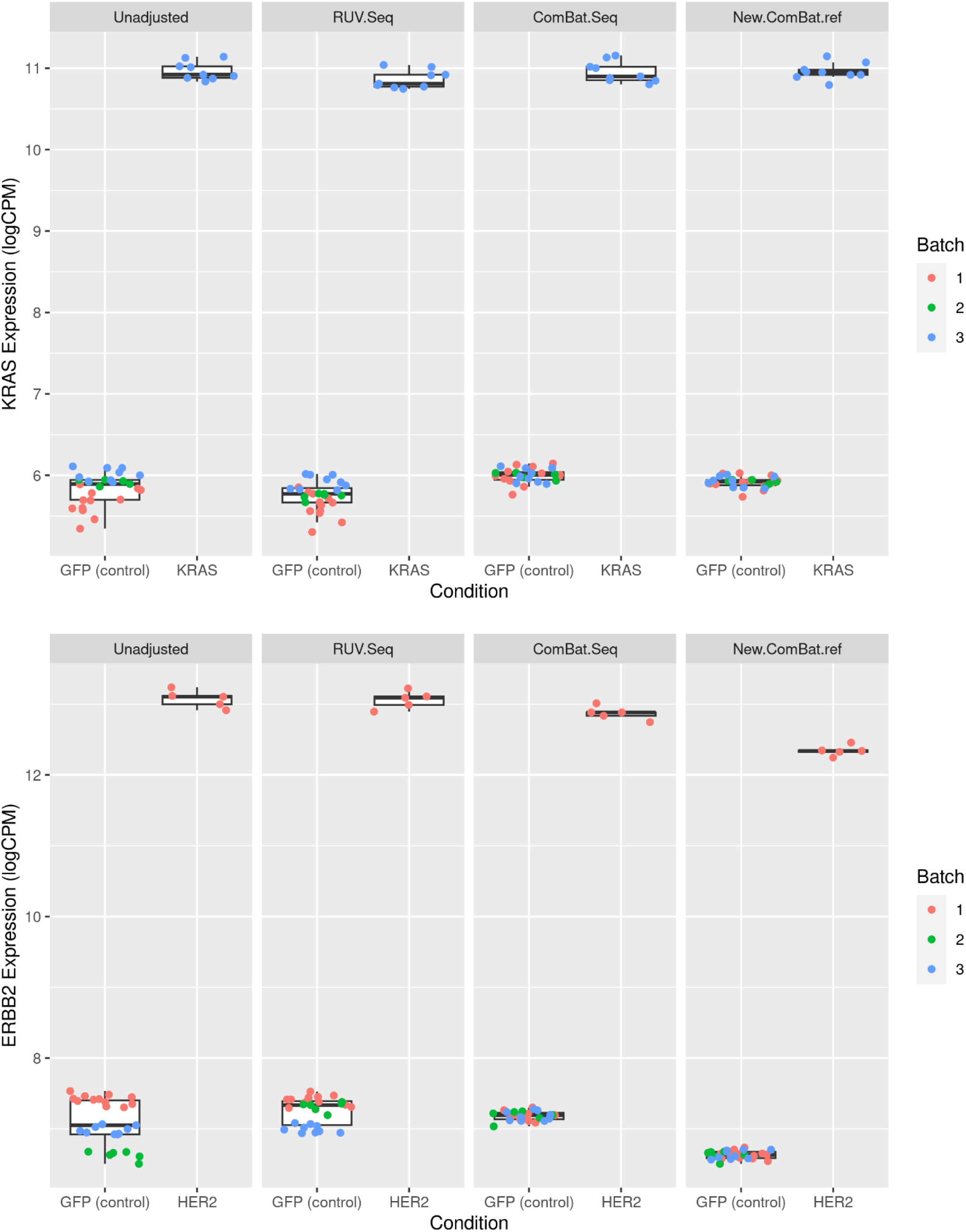
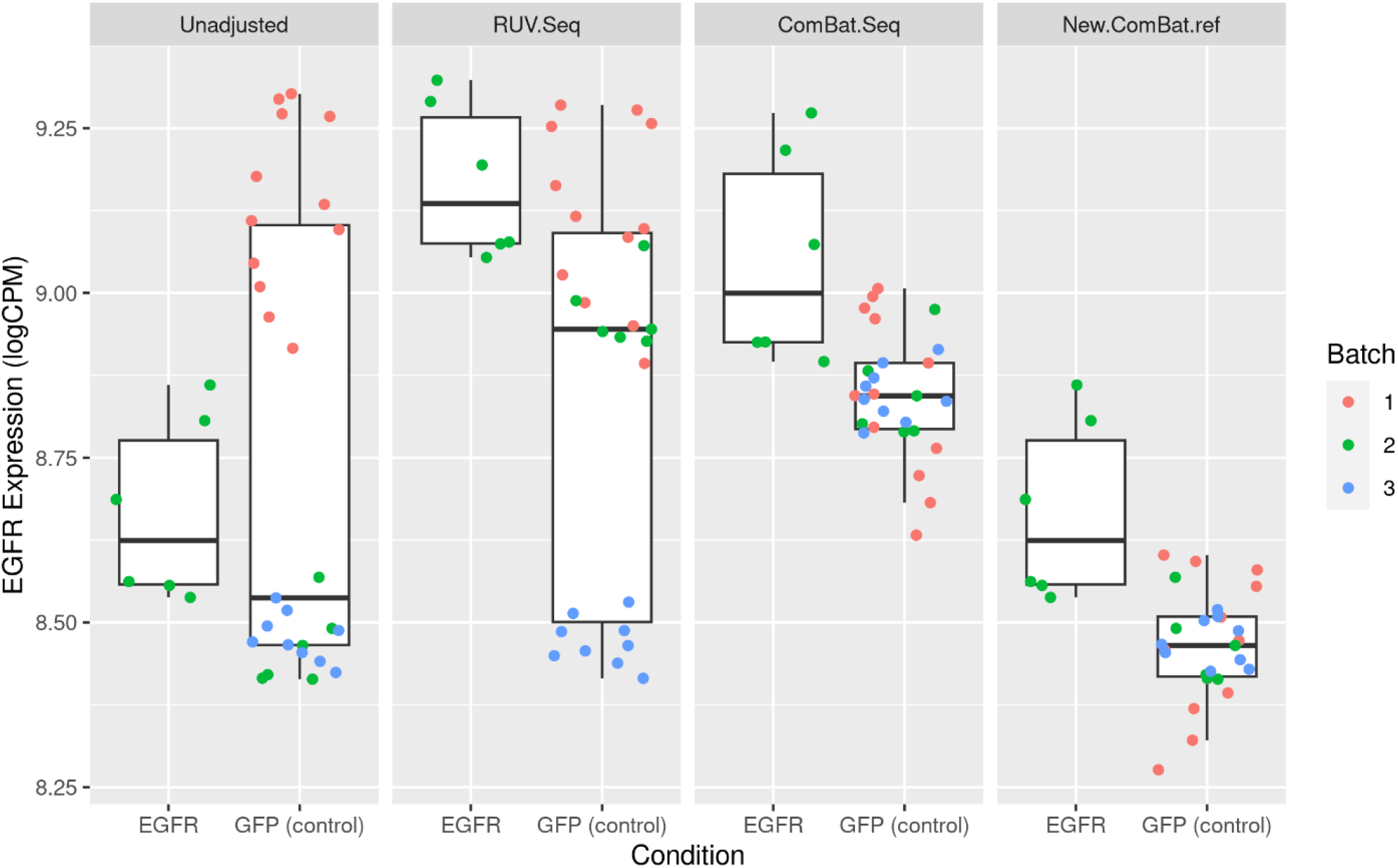
Expressions of KRAS, HER2 and EGFR in unadjusted data and in data adjusted by RUVSeq, ComBat-seq and new ComBat-ref methods. The ComBat-ref adjustment keeps the counts of EGFR genes in the EGFR samples (reference batch) and adjusts its counts in other batches toward the reference. The ComBat-ref result shows more significant differential expression of EGFR gene in the EGFR sample vs. control, verified by the more significant FDR value in DE analysis.

**Supplemental Figure 3.**
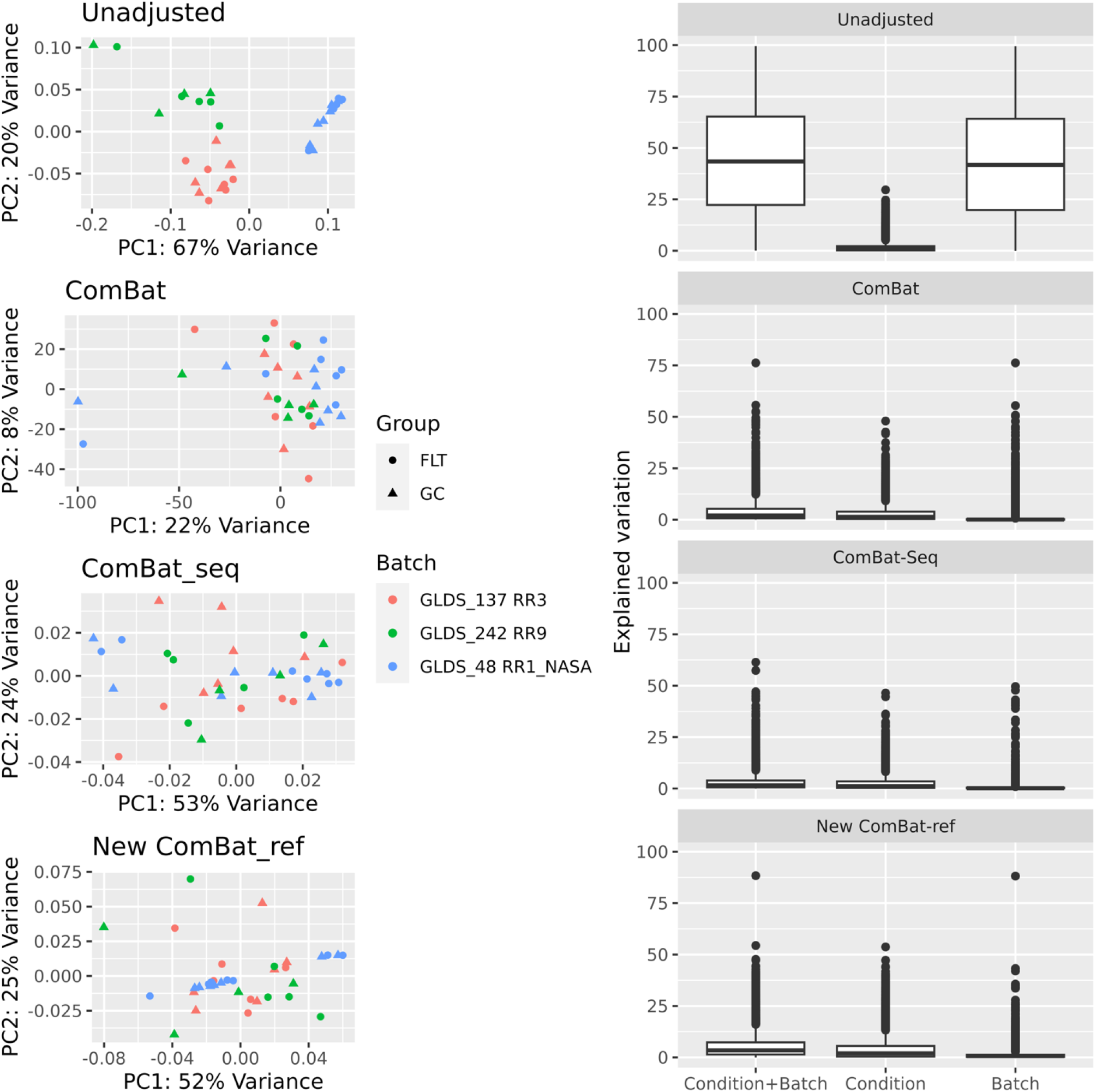
PCA plots of NASA GeneLab dataset for unadjusted data and data adjusted by ComBat, ComBat-seq and the new ComBat-ref method respectively. All three methods are able to remove most batch effects in the original data. ComBat-ref appears to preserve more biological signals in the variations by condition in the explained variation plot on the right.

**Supplemental Table 1.** The number of differentially expressed (DE) genes identified in unadjusted GFRN data and data adjusted using various methods. The DE genes were identified using edgeR with FDR ≤ 0.05. ComBat-ref demonstrates higher power by identifying more DE genes than ComBat-seq in all three samples.

|  | Unadjusted | OneStep | ComBat | RUVseq | SVASEq | ComBat-seq | ComBat-ref |
| --- | --- | --- | --- | --- | --- | --- | --- |
| **her2** | 608 | 3510 | 5916 | 2811 | 1419 | 4727 | 5823 |
| **egfr** | 9798 | 1978 | 4341 | 1330 | 9084 | 4107 | 4934 |
| **kraswt** | 4162 | 3124 | 4809 | 8669 | 5987 | 4991 | 6194 |

**Supplemental Table 2.**
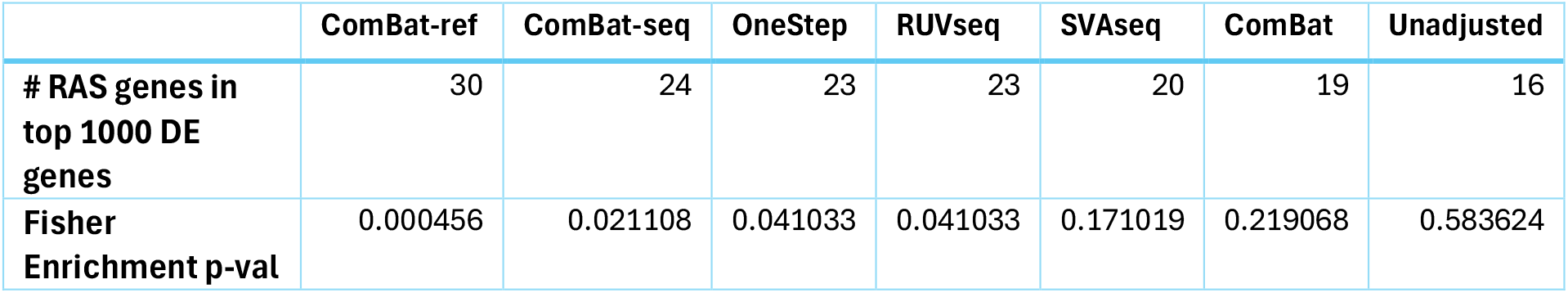
The number of RAS genes found in the top 1000 DE genes identified using various batch adjustment methods. ComBat-ref identified most RAS genes as top DE genes among all methods, with a much more significant p-value for enrichment analysis.

**Supplemental Table 3.** Three batches of NASA GeneLab datasets used in this paper.

| Batch | Mission | Library Preparation | Sequencing Facility |
| --- | --- | --- | --- |
| **GLDS_137** | RR3 | ribodepleted | UCDavis |
| **GLDS_242** | RR9 | ribodepleted | GeneLabSPL |
| **GLDS_48** | RR1_NASA | polyA | UCDavis |

## Notes

https://github.com/xiaoyu12/Combat-ref

### Competing Interest Statement

The authors have declared no competing interest.

